# Contrasting effects of nutrients and consumers on tree colonization and growth during secondary succession

**DOI:** 10.1101/2020.07.13.201129

**Authors:** Robert W. Heckman, Fletcher W. Halliday, Peter A. Wilfahrt

**Affiliations:** Department of Biology, University of North Carolina, Chapel Hill, NC 27599; Environment, Ecology and Energy Program, University of North Carolina, Chapel Hill, NC, 27599; Department of Integrative Biology, University of Texas at Austin, Austin, TX 78712; Department of Evolutionary Biology and Environmental Studies, University of Zurich, Zurich, Switzerland; Department of Ecology, Evolution and Behavior, University of Minnesota, St. Paul, MN 55108

**Keywords:** community assembly, diversity-invasibility, fertilization, insect herbivory, loblolly pine, old fields, *Pinus taeda*, plant pathogens, top-down, bottom-up, tree establishment

## Abstract

For succession to proceed from herbaceous to woody dominance, trees must colonize herbaceous communities and grow. Success across these two phases of succession might result from different interactions with the herbaceous community. First, colonizing trees must compete against larger, established herbs, while subsequent growth occurs among similarly sized or smaller herbs. This shift from colonization to growth may cause three drivers of secondary succession— nutrients, consumers, and herbaceous diversity—to differentially affect tree colonization and growth. Initially, these drivers should favor larger, established herbs, reducing colonization. Later, when established trees can better compete with herbs, these drivers should benefit trees and increase their growth. In a four-year study, we added nutrients to, excluded aboveground consumers from, and manipulated initial richness of, the herbaceous community, then allowed trees to naturally colonize these communities (from intact seedbanks or as seed-rain) and grow. Nutrients and consumers had opposing effects on tree colonization and growth: adding nutrients and excluding consumers reduced tree colonization, but later increased established tree growth (height, basal diameter). Together, this shows stage-specific impacts of nutrients and consumers that may improve predictions of the rate and trajectory of succession: factors that initially limited tree colonization later helped established trees to grow.

## Introduction

For succession to proceed, trees must first colonize, then grow in herbaceous communities (Oosting 1942, Keever 1950, Wright and Fridley 2010, Fridley and Wright 2018). Success in transitioning from colonization to growth may result from changes in the interactions between trees and herbs (HilleRisLambers et al. 2012): colonizing trees must compete against larger, established herbs, while established trees experience limited size-based disadvantages as they continue to grow. These changing interactions may alter the responses of successional systems to other biotic and abiotic factors. For instance, recent work also suggests that soil nutrients and plant consumers (i.e., herbivores and pathogens) may independently and interactively alter early stages of succession (e.g., Fridley and Wright 2012, Meiners et al. 2015, Wilfahrt et al. 2020). Yet, few studies have explored whether nutrients and consumers exert differing impacts on different stages of succession. Importantly, this implies that the same factors that inhibit tree colonization can increase the growth of established trees later in succession, hampering the ability to predict how biotic and abiotic drivers will affect the speed of succession.

Changes in tree-herb interactions between the colonization and growth stages of succession are key for understanding how nutrients and consumers will impact succession. When trees first colonize herbaceous communities, they have a large size-based competitive disadvantage. This interaction may change with soil nutrient availability and consumer pressure (Tilman 2004). High nutrient availability often increases productivity and litter accumulation, reducing light availability (Hautier et al. 2009, Borer et al. 2014, Wilfahrt et al. 2020), and potentially limiting tree colonization (Sarneel et al. 2016). Consumers can reduce the size of resident plant populations and the performance of individuals (Alexander 2010), which may enhance tree colonization. Nutrients and consumers may also jointly alter tree colonization: increased nutrient supply can shift communities toward species and individuals that allocate little to defense against herbivores and pathogens (Hahn and Maron 2016, Heckman et al. 2019), leading to higher rates of herbivory and disease (Veresoglou et al. 2013, Heckman et al. 2016). If consumers are excluded from nutrient-rich habitats, where consumer impacts are highest, the resident community exploit high nutrient supply without experiencing the negative impacts of consumers (Mattson 1980, Heckman et al. 2016); this could drastically reduce tree colonization.

As trees establish, their size-based disadvantage against herbs should decline over time, making competition more symmetric (Schwinning and Weiner 1998), and allowing coexistence when trees and herbs occupy different niches (Chesson 2000). Niche overlap may decline because trees can capture resources unavailable to herbs by developing deeper roots and taller stems and may share few pathogens and herbivores with herbs (Gilbert and Webb 2007, Chesson and Kuang 2008, Craine and Dybzinski 2013). Ultimately, trees will outcompete herbs for light, which is often limiting in nutrient-rich environments (Hautier et al. 2009). As such, factors like high nutrient availability and low consumer pressure, which favored the herbaceous community earlier and slowed succession, could benefit established trees later and accelerate succession.

Nutrient- and consumer-mediated interactions between herbs and trees may also change with herbaceous diversity. More diverse communities often exhibit lower light and soil nutrient availability, less disease and herbivory, and higher productivity (Loreau and Hector 2001, Tilman 2004, Maron et al. 2011, Halliday et al. 2019). This may further enhance the competitive advantage of established herbs over tree seedlings in diverse communities and reduce colonization (Mattingly and Reynolds 2014, Heckman et al. 2017, Wilfahrt et al. 2020). But as trees establish and grow, their competitive disadvantage against herbs should decline, allowing trees and herbs exhibiting sufficiently large niche differences to coexist (Chesson 2000) and reducing the influence of herbaceous diversity on the growth of established trees.

Ultimately, the speed of succession results from the ability of trees to establish and grow within herbaceous communities, which is determined by the abiotic and biotic conditions in the community (Fridley and Wright 2018, Wilfahrt et al. 2020). In this study we examine whether nutrients, consumers, and initial plant community richness interact to influence tree colonization in an old field community. Among established individuals of a dominant early successional tree, we further examine whether these factors interactively influence tree growth.

## Methods

We performed this study at Widener Farm, an old field in Duke Forest Research and Teaching Lab (Orange County, NC, USA) that produced row crops until 1996. Since 1996, the site has been mowed to maintain herbaceous dominance by native species common in North Carolina Piedmont old fields (Oosting 1942) and several exotic species (Heckman et al. 2016).

The study employed a randomized complete block design with three factorial treatments: we manipulated native herbaceous plant richness with multiple community compositions at each level of richness; access by foliar fungal pathogens and insect herbivores; and soil nutrient supply. This yielded a study that comprised 240 plots (5 replicate blocks × 2 nutrient supply levels × 2 consumer access levels × 2 richness levels × 6 native community compositions).

### Plant composition and species richness

In May 2011, we established five spatial blocks, each containing 48 1 m^2^ plots with 1 m aisles. We first applied glyphosate herbicide (Riverdale^®^ Razor^®^ Pro, Nufarm Americas Inc, Burr Ridge, IL) to each plot, removed all dead vegetation, and covered plots with landscape fabric, while avoiding disturbance to the existing seed bank. We assigned each plot to one of two richness levels, monoculture or five-species polyculture. From a pool of six native herbaceous perennial species, we assembled twelve planted communities: six monocultures and six five-species polycultures, where one species was excluded from each community. All species were already present locally, and included three grasses—*Andropogon virginicus, Setaria parviflora, Tridens flavus*, and three forbs—*Packera anonyma, Scutellaria integrifolia, Solidago pinetorum*.

We propagated all species in the greenhouse at the University of North Carolina at Chapel Hill. Each species was planted between June and September 2011 in 1-2 days when seedlings were large enough to survive transplant stress. In early summer 2012, we replaced all individuals that had died. To minimize recruitment from the seedbank while establishing the species richness treatment, seedlings were planted into small holes in the landscape fabric covering the plot. Plots contained 41 individuals, each spaced ∼10 cm from its nearest neighbors in a checkerboard pattern. Polycultures contained 9 individuals of one randomly chosen species and 8 individuals of the other 4 species. In July 2012, we removed landscape fabric from plots and weeded non-planted individuals by hand. We then allowed natural colonization for the duration of the study. Thus, the species richness treatments represent initial conditions.

### Nutrient supply and consumer access treatments

We began consumer access and nutrient supply treatments in July 2012. To manipulate access by foliar fungal pathogens and insect herbivores, each plot was assigned to one of two treatments (sprayed with fungicide and insecticide vs. not sprayed). From July 2012 through September 2015, we sprayed non-systemic broad-spectrum biocides on all aboveground biomass every two to three weeks during the main growing season (April-October). Neither the fungicide (mancozeb, Dithane^®^ DF, Dow AgroSciences, Indianapolis, IN) nor the insecticide (es-fenvalerate, Asana^®^ XL, Dupont, Wilmington, DE) had any non-target effects on plant growth under greenhouse conditions (Heckman et al. 2016). Similarly, this fungicide has no adverse effects on mycorrhizal fungi when used as recommended (Parker and Gilbert 2007). Together, these biocides reduced foliar damage in this study by >55% (Heckman et al. 2017).

To manipulate soil nutrient supply, each plot was assigned to one of two treatments (fertilized with 10 g N m^-2^ yr^-1^ as slow-release urea, 10 g P m^-2^ yr^-1^ as triple super phosphate, and 10 g K m^-2^ yr^-1^ as potassium sulphate vs. not fertilized). This level of fertilization has been shown to alleviate limitation by N, P, and K across a range of grassland habitats (Fay et al. 2015). In 2012, we fertilized plots in July, and in subsequent years, we fertilized in early May.

### Tree colonization and growth

To examine tree colonization, we identified all plant species in a marked 0.75 × 0.75 m subplot in the center of each plot in September 2012 – 2015 (Wilfahrt et al. 2020); 11 tree species had colonized one or more plots (*Acer rubrum, Celtis laevigata, Cercis canadensis, Cornus florida, Fraxinus* sp., *Juniperus virginiana, Liquidambar styraciflua, Liriodendron tulipifera, Pinus taeda, Sassafras albidum*, and *Ulmus alata*). To examine the growth of established trees, in May 2016, we measured the height and basal diameter of each *P. taeda* individual. We focused on *P. taeda* because it is the most abundant early successional tree in the region and within this study (Oosting 1942, Wright and Fridley 2010).

### Data analysis

We took two approaches to examine the effects of treatments on tree dynamics: evaluating the presence of trees across all plots and evaluating tree presence and performance only in plots containing trees. Modeling tree presence in all plots allowed us to understand overall treatment effects, but did not account for stochasticity in natural seed rain. In contrast, modeling tree growth and colonization time only in plots containing trees allowed us to evaluate treatment effects while greatly reducing stochasticity in natural seed rain. We analyzed all data in R version 3.5.3 (R Foundation for Statistical Computing, Vienna 2019).

To model the independent and interactive effects of initial richness, fertilization, and spraying on tree colonization across all plots (i.e., the presence of trees in a plot), we used the glmmTMB package for generalized linear mixed models (Brooks et al. 2017) with binomial errors and a logit link. This model also included year of observation as a continuous fixed effect, which could interact with treatment effects. We assessed model significance with Wald tests using the ‘Anova’ function in the car package (Fox and Weisberg 2018). To evaluate how initial richness, fertilization, and spraying independently and interactively influenced tree growth and colonization time in plots containing trees, we analyzed three responses using the ‘lme’ function in the nlme package (Pinheiro et al. 2016): the height and basal diameter of *P. taeda* in spring 2016 and the earliest year (fall 2012-2015) in which trees had colonized a plot. In all models, fertilization, spraying, initial richness, and their interactions were categorical fixed effects.

Following Schmid et al. (2002), planted community composition was a random effect; plot was nested within composition to account for repeated sampling in the tree colonization GLMM. We simplified fixed effects following Zuur et al. (2009).

## Results

As expected in an old field undergoing succession, the presence of trees increased over time (Time: P = 0.027; Table S1) and this effect was interactively altered by nutrients and consumers (Nutrients × Consumers × Time: P < 0.001; Table S1; Figure 1). Spraying and fertilization each reduced the rate at which trees colonized plots relative to controls (P < 0.001 for each contrast), but spraying did not significantly change the colonization rate in fertilized plots (P = 0.07). Among plots containing trees, spraying and fertilization also interactively altered the timing of establishment (Nutrients × Consumers: P = 0.02; Table S2; Figure S1). Spraying advanced the average colonization date by 0.91 years in unfertilized plots (P = 0.01), but did not affect colonization time in fertilized plots (P = 0.91). Similarly, fertilization advanced the average colonization date by 0.92 years in unsprayed plots (P = 0.018), but did not affect colonization time in sprayed plots (P = 0.91). Initial richness did not influence either response (Table S1; Table S2). This indicates that trees colonized sprayed and fertilized plots less frequently, but trees that colonized sprayed and fertilized plots did so earlier.

**Figure 1.**
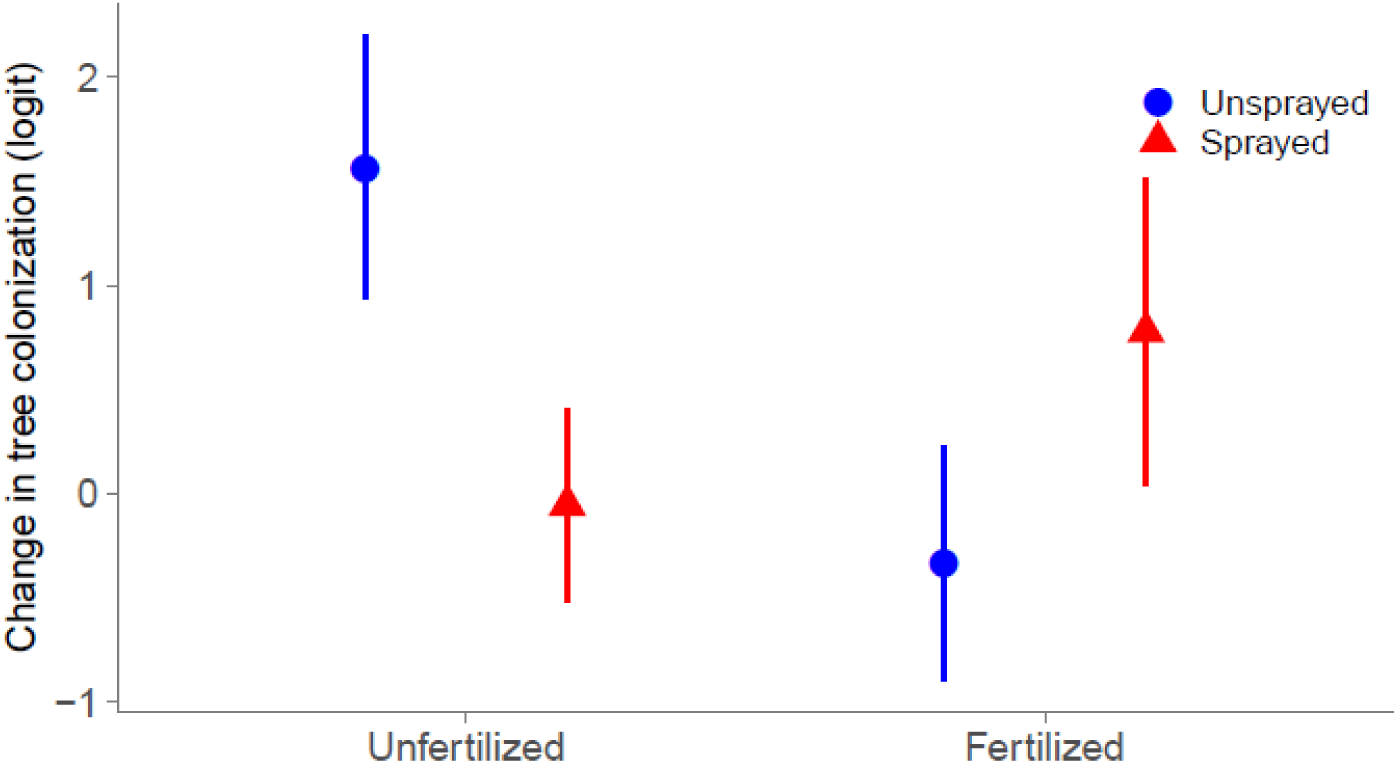
Effects of nutrient supply (unfertilized, fertilized with NPK) and consumer access (unsprayed, sprayed with aboveground fungicide and insecticide) on logit change in colonization of plots (presence/absence) by all tree species between the beginning of the study in 2012 and 2015 (N = 240 plots; mean ± 95% confidence intervals), calculated using a generalized linear mixed model with binomial errors and a logit link. Error bars overlapping 0 denote no change in the rate at which trees colonized plots over the course of the study; positive values indicate that the rate at which trees colonized plots increased over the course of the study

Whereas fertilization and spraying reduced tree colonization, these treatments had the opposite effect on growth of the focal species, *P. taeda*. Specifically, after four years of treatments, nutrients and consumers interactively altered two measures of *P. taeda* growth: basal diameter and height (Basal diameter, Nutrients × Consumers: P = 0.002; Height, Nutrients × Consumers: P = 0.002; Table S3; Figure 2a, 2b). Spraying increased basal diameter by 105% and height by 81% in unfertilized plots (Basal diameter: P = 0.002; Height: P = 0.001), but not in fertilized plots (Basal diameter: P = 0.60; Height: P = 0.67), while fertilization increased basal diameter by 84% and height by 55% in unsprayed plots (Basal diameter: P = 0.031; Height: P = 0.055), but not in sprayed plots (Basal diameter: P = 0.21; Height: P = 0.13). Similar to tree establishment, initial richness had no effect on *P. taeda* growth (Table S3).

**Figure 2.**
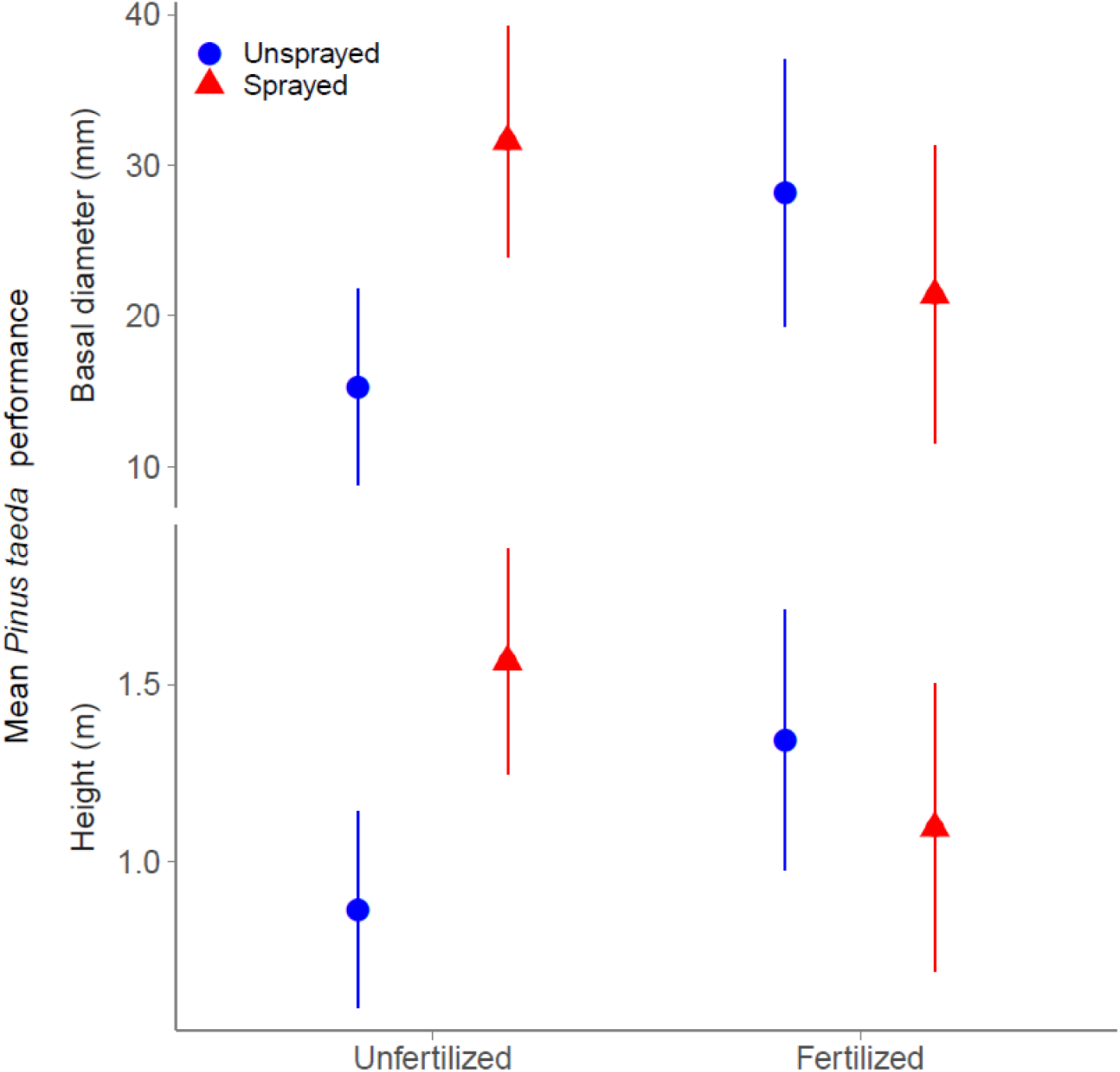
Effects of nutrient supply (unfertilized, fertilized with NPK) and consumer access (unsprayed, sprayed with aboveground fungicide and insecticide) on performance of *Pinus taeda* (basal diameter and height) in spring 2016 (N = 63 plots; mean ± 95% confidence intervals), after four growing seasons of experimental treatments, calculated using linear mixed models with restricted maximum likelihood estimation

Together, these results reveal contrasting effects of nutrients and consumers on tree colonization and growth. Fertilization and spraying hindered tree colonization after the first year of the study, possibly because these treatments disproportionately benefitted the herbaceous community. But when trees established, fertilization and spraying enhanced their growth.

## Discussion

These results provide evidence for contrasting effects of nutrients and consumers on two early stages of secondary succession, tree colonization and growth. These contrasting effects are likely driven by a change in the role of the herbaceous community during succession (Pickett et al. 1987, Meiners et al. 2015): herbaceous residents can be critical for hindering colonization and establishment (Smit and Olff 1998, Rebele 2013), but established trees might outcompete, or avoid competing with, herbs. Thus, factors like high nutrient supply and low consumer pressure largely prevented tree colonization by benefiting the herbaceous community (Sarneel et al. 2016, Heckman et al. 2017), while later promoting growth among the trees that did establish.

Contrasting effects of competitors, consumers, and nutrients on colonization and growth could help explain some of the conflicting results seen in earlier studies of the drivers of secondary succession (e.g., Gill and Marks 1991, Rebele 2013, Fridley and Wright 2018).

As predicted, nutrient addition and consumer exclusion each reduced tree colonization, potentially by several mechanisms. Adding nutrients and excluding consumers reduced light availability (Wilfahrt et al. 2020), which was a key driver of herbaceous colonization in another study at this site (R.W. Heckman *unpublished data*) and for community assembly broadly (Hautier et al. 2009, Harpole et al. 2017). Moreover, because herbs were at a higher density than colonizing trees, they likely experienced stronger negative consumer impacts through high density-dependent consumer pressure (Chesson and Kuang 2008, Mordecai 2011). Thus, excluding consumers likely reduced tree colonization by favoring the herbaceous community.

Together, this suggests that the larger size and higher density of herbs relative to trees allowed them to exploit the more favorable conditions of adding nutrients and excluding consumers.

Whereas adding nutrients and excluding consumers reduced tree colonization, these factors increased the growth of established trees, which we measured using the focal tree species, *P. taeda*. Several processes likely contributed to this effect. First, *P. taeda* individuals could occupy a niche distinct enough from the herbaceous community to largely avoid competition (Chesson 2000). This could have occurred because *P. taeda* is phylogenetically and functionally distinct from the herbaceous community (Mayfield and Levine 2010). By exploiting this niche difference, *P. taeda* would benefit from increased nutrient supply and release from consumers without experiencing increased competition intensity (Chesson and Kuang 2008). Second, once *P. taeda* established, and reached the herbaceous canopy, these trees may have been released from competition for light and become more apparent to plant consumers, resulting in stronger competition for soil nutrients and stronger regulation by consumers (Schwinning and Weiner 1998, Chesson and Kuang 2008, Mordecai 2011). Finally, *P. taeda* may have outcompeted the herbaceous community. Conditions promoting strong competition for light, like nutrient addition, favor tall *P. taeda* individuals over short herbs (Hautier et al. 2009, Craine and Dybzinski 2013). Importantly, even if *P. taeda* successfully exploited a niche difference, its superior ability to compete for light will, in the absence of disturbance, drive its herbaceous competitors to low abundance or local extinction (Craine and Dybzinski 2013).

This study demonstrates the importance of nutrients and consumers in driving early succession but has several limitations. First, we did not measure damage on *P. taeda* or any other tree in this study. However, past research has shown that our spraying approach effectively reduces damage to numerous species without having biotoxic or biostimulatory effects (Heckman et al. 2016, Heckman et al. 2017). Second, our five-year study did not cover the entire, decades-long duration of succession. However, succession in southern US old fields proceeds rapidly (Oosting 1942, Keever 1950, Fridley and Wright 2018, Wilfahrt et al. 2020), and our study captured the critical early stages of old field succession, which determine the trajectory of forest development (Fridley and Wright 2018). Finally, because the field surrounding this experiment was maintained by mowing, very few trees in the field were large enough to produce seeds. Consequently, most trees colonized plots from the forest edge or resident seedbank, resulting in stochastic colonization and establishment—by the end of the study, even *P. taeda*, the most successful early successional tree, had only established in ∼ 25% of plots. Future experiments could overcome this limitation by adding seeds or seedlings.

Our results suggest that competition with resident herbs is an important driver of tree colonization and growth during secondary succession; yet, counterintuitively, neither tree colonization nor growth were affected by initial richness of the herbaceous community. This raises an important question: if competition with resident herbs was so important for tree colonization or growth, why were no effects of initial herbaceous richness detected? One possible explanation for this result is that increasing niche complementarity, which is often associated with reduced invasion or increased stability in more diverse communities, was not necessary to inhibit colonization or impact tree growth (Shea and Chesson 2002, Seabloom 2007). This may occur because trees and herbs were competing primarily for an asymmetric resource, light. In old fields, where most species are shade intolerant, there is limited opportunity to partition light in a way that promotes coexistence. Rather, when vegetation is dense enough to create a closed canopy, taller individuals gain a considerable competitive advantage irrespective of niche differentiation (Westoby 1998). Moreover, because we did not maintain richness treatments beyond July 2012, any richness effect declined over time (Halliday et al. 2019, Wilfahrt et al. 2020).

Our results indicate that the same drivers can affect each stage of early succession differently, potentially resolving idiosyncrasies in previous studies. Within a site, many studies have shown that nutrient supply and consumer pressure are important drivers of succession (Pickett et al. 1987, Meiners et al. 2015), but the strength and direction of these effects often differed (Gill and Marks 1991, Rebele 2013). Many of these results are seen as supporting different models of succession. For instance, the succession models of Connell and Slatyer (1977) describe three possible outcomes: inhibition, tolerance, and facilitation. In our study, the earliest stages of succession—when the herbaceous community prevented tree colonization— were consistent with the inhibition model. Later, when established trees responded positively to nutrient addition and consumer exclusion, herbs and trees may have exhibited more neutral interactions, consistent with the tolerance model. However, when studies do not account for changing interactions between herbs and trees, these differences may be obscured. Thus, discrepancies in the importance of nutrients and consumers among past studies may have resulted from testing colonization and growth together instead of considering each separately.

### Conclusions

In this study, two early stages of succession from herbaceous to woody dominance were influenced by nutrients and consumers in contrasting ways, suggesting an important shift in the ecological drivers of secondary succession. Nutrient addition and consumer exclusion limited tree colonization, likely through an indirect route mediated by the herbaceous community. Unlike establishment, growth of the focal tree, *P. taeda* increased with nutrient addition and consumer exclusion, perhaps because trees overcame competition for light with the herbaceous community. Thus, these two factors, which are so critical to many aspects of community ecology (HilleRisLambers et al. 2012), had contrasting effects on different stages of succession. As large tracts of former farmland are abandoned and undergo secondary succession (Wright and Fridley 2010), the speed of this process, and the carbon that it sequesters, may be determined to a large extent by consumers and nutrient availability.

## Acknowledgments

We thank C. Mitchell, J. Bruno, J. Umbanhowar J. Wright, and members of the Mitchell lab provided helpful suggestions for the design of this experiment. Many members of the Plant Ecology Lab, Mitchell Lab, and Duke Forest provided field assistance and botanical expertise. This study was funded by a Doctoral Dissertation Improvement Grant to RWH (NSF-DEB-1311289). RWH and PAW were supported by UNC’s Alma Holland Beers Scholarship and WC Coker Fellowship. RWH and FWH were supported by the UNC Graduate School Dissertation Completion Fellowship. FWH was supported by the NSF Graduate Research Fellowship.

## Author contributions

RWH, FWH, and PAW conceived and implemented the experiment. RWH analyzed the data and wrote the first draft. All authors contributed to revisions of the manuscript.

## Data accessibility

Upon acceptance, data will be archived in the Dryad Digital Repository.

