## Supplementary Materials for "Contrasting effects of nutrients and consumers on tree colonization and growth during secondary succession"

The following Supporting Information is available for this article:

**Table S1** Summary of generalized linear mixed model results, tree colonization

**Table S2** Summary of linear mixed model results, year of tree colonization

**Table S3** Summary of linear mixed model results, tree performance

**Figure S1** Effects of nutrient supply and consumer access on initial year of tree colonization

**Table S1** ANOVA table showing effects of treatments on colonization of plots (presence/absence) by all tree species between the beginning of the study in 2012 and 2015 (N = 240 plots)

| Parameter | $\chi^2$ | P value |
|---|---:|---:|
| Consumers | 0.607 | 0.436 |
| Nutrients | 2.397 | 0.122 |
| Richness | 0.071 | 0.79 |
| Year | 4.92 | 0.027 |
| Consumers × Nutrients | 0.123 | 0.726 |
| Consumers × Year | 2.583 | 0.108 |
| Nutrients × Year | 3.389 | 0.066 |
| Richness × Year | 0.539 | 0.463 |
| Consumers × Nutrients × Year | 18.285 | < 0.001 |

**Table S2** ANOVA table showing effects of treatments on initial year of tree colonization in plots containing trees (N = 75 plots)

| Parameter | F value | P value |
|---|---:|---:|
| Intercept | 29.036 | < 0.001 |
| Consumers | 9.755 | 0.003 |
| Nutrients | 8.481 | 0.005 |
| Richness | 0.038 | 0.849 |
| Consumers × Nutrients | 5.685 | 0.02 |

**Table S3** ANOVA table showing effects of treatments on performance of *Pinus taeda* (basal diameter and height) in spring 2016 (N = 63 plots), after four growing seasons of experimental treatments

| Response | Parameter | F value | P value |
|---|---|---:|---:|
| Basal diameter | Intercept | 12.621 | 0.001 |
| | Consumers | 13.851 | 0.001 |
| | Nutrients | 7.402 | 0.009 |
| | Richness | 0.343 | 0.571 |
| | Consumers × Nutrients | 10.504 | 0.002 |
| Height | Intercept | 20.946 | < 0.001 |
| | Consumers | 15.322 | < 0.001 |
| | Nutrients | 6.244 | 0.016 |
| | Richness | 1.014 | 0.338 |
| | Consumers × Nutrients | 10.773 | 0.002 |

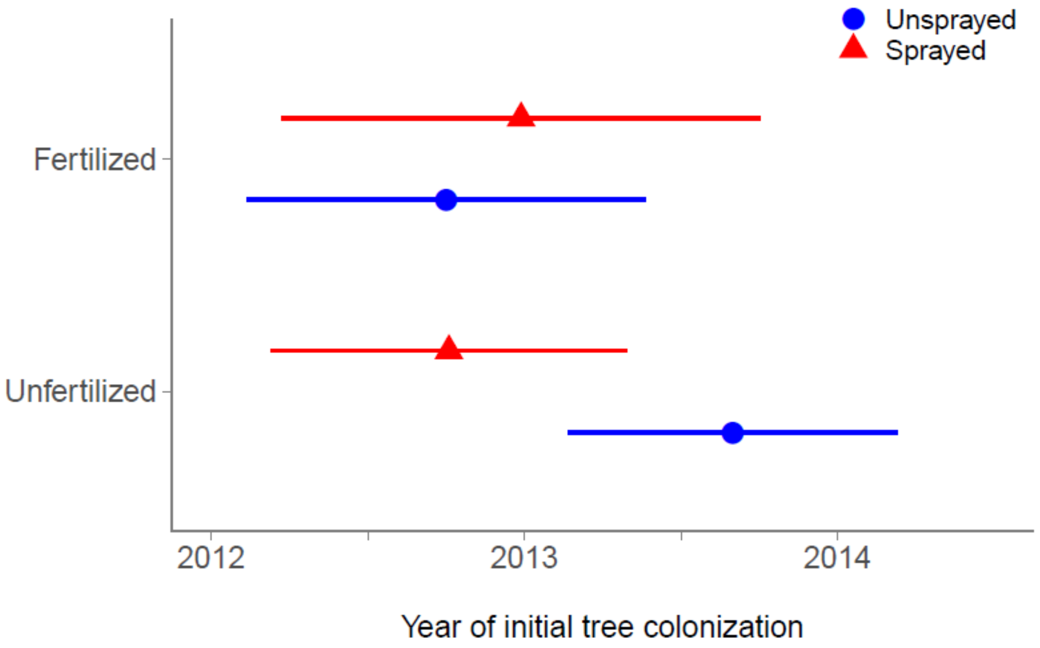


**Figure S1** Effects of nutrient supply (unfertilized, fertilized with NPK) and consumer access (unsprayed, sprayed with aboveground fungicide and insecticide) on year of initial tree colonization of plots containing trees (N = 75 plots; mean ± 95% confidence intervals) calculated using linear mixed models with restricted maximum likelihood estimation. Experimental treatments began in 2012 and continued through 2015
